# Astrocytes integrate local sensory and brain-wide neuromodulatory signals

**DOI:** 10.1101/381434

**Authors:** Michal Slezak, Steffen Kandler, Paul P. Van Veldhoven, Vincent Bonin, Matthew G. Holt

**Author notes:** These authors contributed equally to the work (Michal Slezak, Steffen Kandler). Co-senior authors (Vincent Bonin, Matthew G. Holt). Current Address: BioMedX GmbH, BioMedX Innovation Center, Im Neuenheimer Feld 515, D69120 Heidelberg, Germany (Michal Slezak). **Correspondence** Dr. Matthew Holt, Laboratory of Glia Biology, VIB-KU Leuven Center for Brain and Disease Research, KU Leuven Department of Neuroscience, O&N IV, Herestraat 49 - box 602, Leuven 3000, Belgium, Dr. Vincent Bonin, IMEC NERF, Kapeldreef 75, Leuven 3001, Belgium.

## Abstract

Astrocytes play multiple functions in the central nervous system, from control of blood flow through to modulation of synaptic activity. Transient increases in intracellular Ca^2+^ are thought to control these activities. The prevailing concept is that these Ca^2+^ transients are triggered by distinct pathways, with little mechanistic and functional overlap. Here we demonstrate that astrocytes in visual cortex of mice encode local visual signals in conjunction with arousal state, functioning as multi-modal integrators. Such activity adds an additional layer of complexity to astrocyte function and may enable astrocytes to specifically and subtly regulate local network activity and plasticity.

---

Astrocytes are one of the major cell types in the central nervous system (CNS), interacting with neurons, other glia and blood vessels^1^. This allows them to modulate neural activity and behavior^2^, promote myelination^3^ and control blood flow^4^, often in a co-ordinated fashion. Astrocyte activation is commonly associated with a transient rise in intracellular Ca^2+^, which has been linked correlatively to functional outputs^5, 6^. However, it is becoming increasingly obvious that astrocyte Ca^2+^ signals do not reflect a single mechanism of activation, but rather are involved across a wide range of molecular signaling events underlying the physiological actions of astrocytes^5^. An important, unresolved question is the degree to which those signaling events interact in astrocytes *in vivo*.

Recent studies suggest a path to study interactions across Ca^2+^ signaling pathways in astrocytes. In mice, astrocytes show Ca^2+^ transients in response to motor activity^7, 8^ that are believed to involve the entire cortical astrocyte network^7^ and may be mediated by neuromodulators^7, 9^. Local sensory responses have been described in anesthetized ferrets^6^, but in mice responses are thought to be weak and unreliable^7, 10, 11^. We used GCaMP6 labeling with widefield 1-photon and 2-photon microscopy to investigate astrocyte Ca^2+^ activity in primary visual cortex (V1), in response to visual stimulation, in mice engaged in voluntary locomotor behavior (Figure 1a, b). Consistent with earlier reports^7–9^, locomotion alone induced synchronous increases in astrocyte Ca^2+^ across the posterior cortex (Figure 1c, d, Supplementary Figure 1a, Supplementary Movie 1). In contrast, use of drifting visual stimuli^12^, concomitant to locomotion, elicited waves of activity that travelled retinotopically across the cortex, as the stimulus moved across the visual field (Figure 1e-g, Supplementary Figure 1a, b, Supplementary Movies 2, 3). The direction and speed of the drifting stimuli were directly reflected in the direction and speed at which the responses moved across the cortex (Figure 1e-g). Furthermore, bar speed directly influenced the amplitude of responses and response probability (Supplementary Figure 1c, d), yet the overall time course of the visually-evoked Ca^2+^ transients were invariant to bar speed (Supplementary Figure 1e). This was true both in the widefield images of global astrocyte-related Ca^2^ activity and 2-photon Ca^2+^ responses of individual astrocytes. This suggests that astrocyte responses to visual stimulation, in the mouse cortex, follow local synaptic activity and are triggered in each individual cell, rather than Ca^2+^ spreading through a gap junction-coupled network^13^.

**Figure 1.**
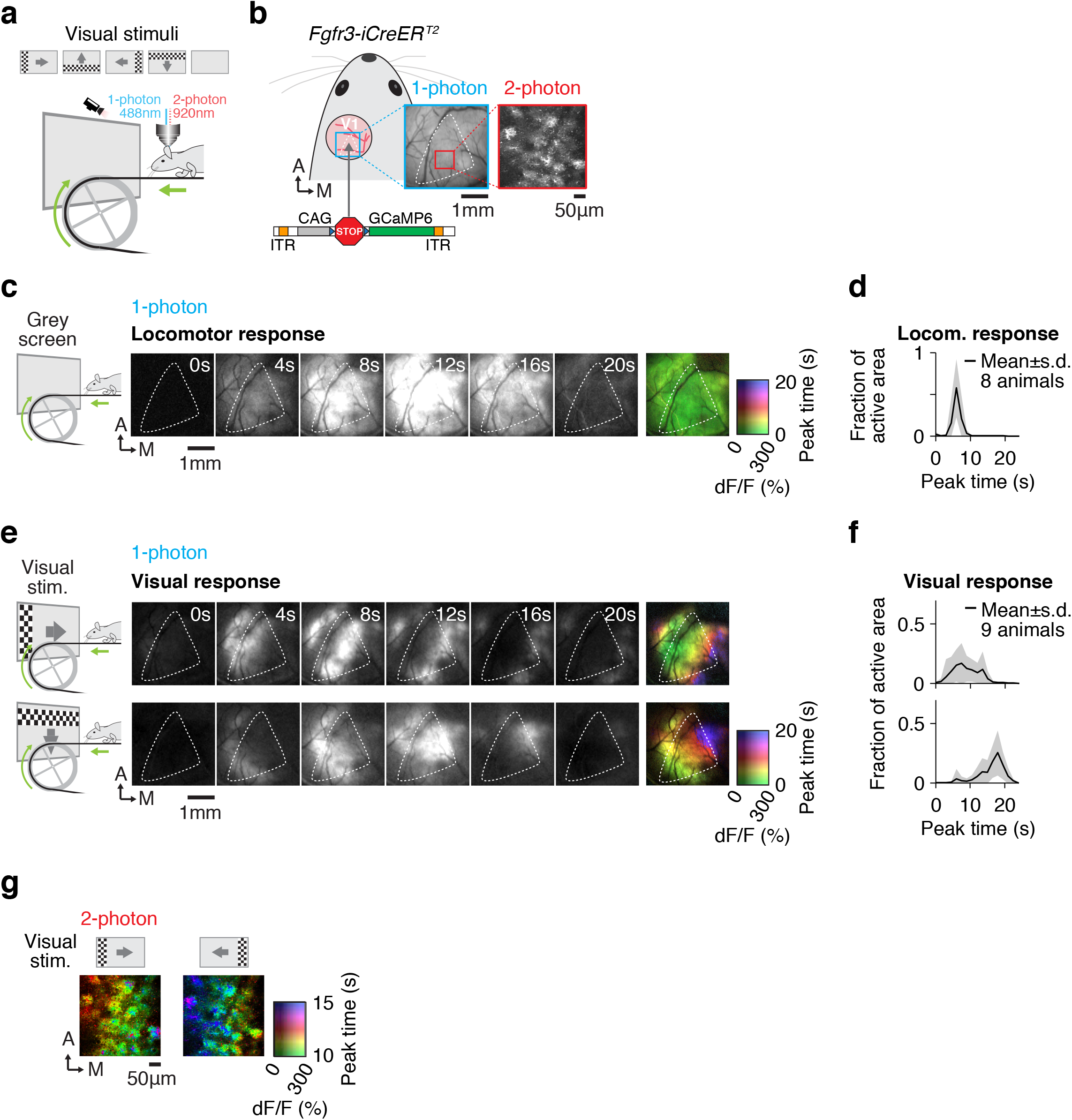
Visually evoked Ca^2+^ increases in astrocytes of the primary visual cortex in mice show retinotopic organization. (a) Mice were head fixed under a multiphoton microscope. Visual stimuli comprised flickering checkerboard bars moving in the 4 cardinal directions and were presented to the right eye of each animal. This eye was continuously monitored for changes in position and pupil size. Animals were free to move on a treadmill during the experiment. (b) *Fgfr3-iCreER^T2^* mice were implanted with a 5 mm cranial window centered over the left visual cortex. GCaMP6 was expressed specifically in astrocytes following stereotactic injection of a flexed AAV construct and administration of tamoxifen. GCaMP6 fluorescence was recorded from extended or local (cellular level) V1 regions of interest (ROIs) using 1-photon (blue ROI) or 2-photon (red ROI) imaging. Anterior (A) and Medial (M). (c, d) 1-photon imaging during bouts of locomotion revealed synchronous increases in GCaMP6 fluorescence across the ROI. Individual black and white frames are taken from Supplementary Movie 1. Numbers indicate the time from onset of locomotion. The pseudo-colored image shows the average time to maximal increase in GCaMP6 fluorescence after locomotion onset. Map is the average of 22 trials from 1 animal. Dashed outlines indicate borders of V1 (c). Quantification of changes in GCaMP6 fluorescence after locomotion onset. Curves indicate mean ± s.d. across animals (d). (e, f) 1-photon imaging during presentation of visual stimuli, with concomitant locomotion, revealed a retinotopic organization of the GCaMP6 response. Direction of stimulus movement is indicated. Individual black and white frames are from Supplementary Movie 2. Numbers indicate the time from onset of visual stimulation. The pseudo-colored image shows the average time to maximal increase in GCaMP6 fluorescence after onset of visual stimulation. Map is the average of 10 trials per stimulus direction in 1 animal. Dashed outlines indicate borders of V1 (e). Quantification of changes in GCaMP6 fluorescence after onset of visual stimulation. Curves indicate mean ± s.d. across animals (f). (g) Pseudo-color, pixel-wise maps of trial averages for vision-induced increases in GCaMP6 fluorescence obtained using 2-photon imaging. Map is the average of 10 trials per stimulus direction in 1 animal.

Next, we examined the possible interplay between visual responses and behavioral state. As described above, clear responses to visual stimuli were discernible during locomotion, but not during stationary epochs (Figure 2a, b, Supplementary Figure 2). Activity during stationary epochs was only mildly different from baseline (Figure 2b). However, amplitudes of visual responses observed during locomotion were larger than responses to locomotion onset alone (Figure 2c). Similar results were obtained for widefield imaging and 2-photon recordings of individual astrocytes (Figure 2b, c, left vs. right). The degree of motor activity did not determine the strength of visual responses (Figure 2d; correlation coefficient = 0.09, p-value = 0.15). However, response amplitude did highly correlate with pupil size (Figure 2e; correlation coefficient = 0.57, p-value ≤ 0.001). Pupil size is often used as a proxy for arousal^14^. This suggests that astrocytes integrate specific visual signals with behavioral state, as sensory responses are effectively gated by arousal.

**Figure 2.**
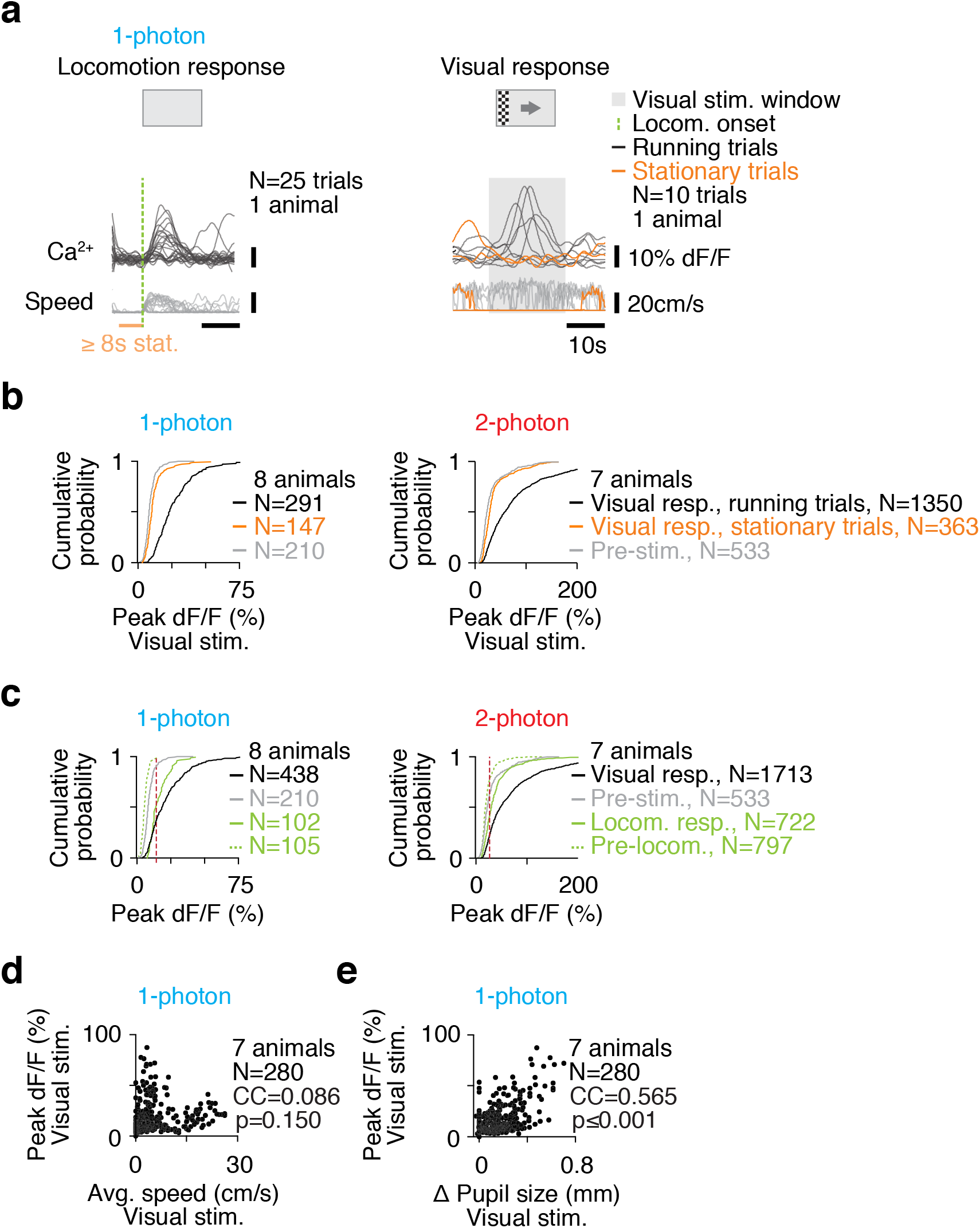
Ca^2+^ signals elicited in visual cortex astrocytes integrate information on sensory input and behavioral state of the mouse. (a, b) Astrocyte responses to visual stimulation are conditional on locomotion. Recordings of Ca^2+^ transients induced by locomotion onset (left), or visual stimulation simultaneous with spontaneous locomotion (right). Dashed line indicates the onset of locomotion (after a stationary period lasting at least 8 s). Orange curves show trials in which the mouse is stationary, showing no movement from 1s before presentation of the visual stimulation and throughout the presentation period. Grey box indicates the time at which visual stimulation was present (a). Cumulative distribution plots showing the peak fluorescence change when the animal is stationary or running during presentation of visual stimuli. Data were recorded in 1-photon (left) or 2-photon mode (right) (b). (c) Visual stimulation during locomotion (black curves) produced a greater response than locomotion alone (green curves). Data were recorded in 1-photon (left) or 2-photon mode (right). Red line indicates the standard deviation threshold. (d, e) Visually-induced responses are correlated with arousal but not locomotion. Scatter plots showing the peak increase in GCaMP6 fluorescence recorded during visual stimulation with voluntary locomotion plotted against the average speed of locomotion (d), or the changes in pupil size that occurred after the onset of stimulation (e). CC, correlation coefficient. N = number of events.

Changes in arousal are linked to release of the neuromodulator noradrenaline, from projections of the locus coeruleus (LC). Noradrenaline is known to trigger Ca^2+^ transients in astrocytes^7, 9^ through activation of the astrocyte-specific α1-adrenoreceptor^15^. To test the contribution of noradrenergic inputs to visual responses, we systemically administered DSP-4^16^ to mice: this toxin leads to selective degeneration of noradrenergic neurons in the cortex and a reduction of noradrenaline levels (Figure 3a). Administration of DSP-4 abolished Ca^2+^ responses to locomotion onset (Figure 3b, c, f), consistent with previous reports^7, 9^. In comparison, responses to visual stimuli were much less impacted, showing primarily a reduction in response amplitude (Figure 3d, g, Supplementary Figure 3a). Crucially, the retinotopic organization of the activity was unaffected by DSP-4 administration (Figure 3e; Supplementary Figure 3b, Supplementary Movie 4), suggesting it is not defined by noradrenaline.

**Figure 3.**
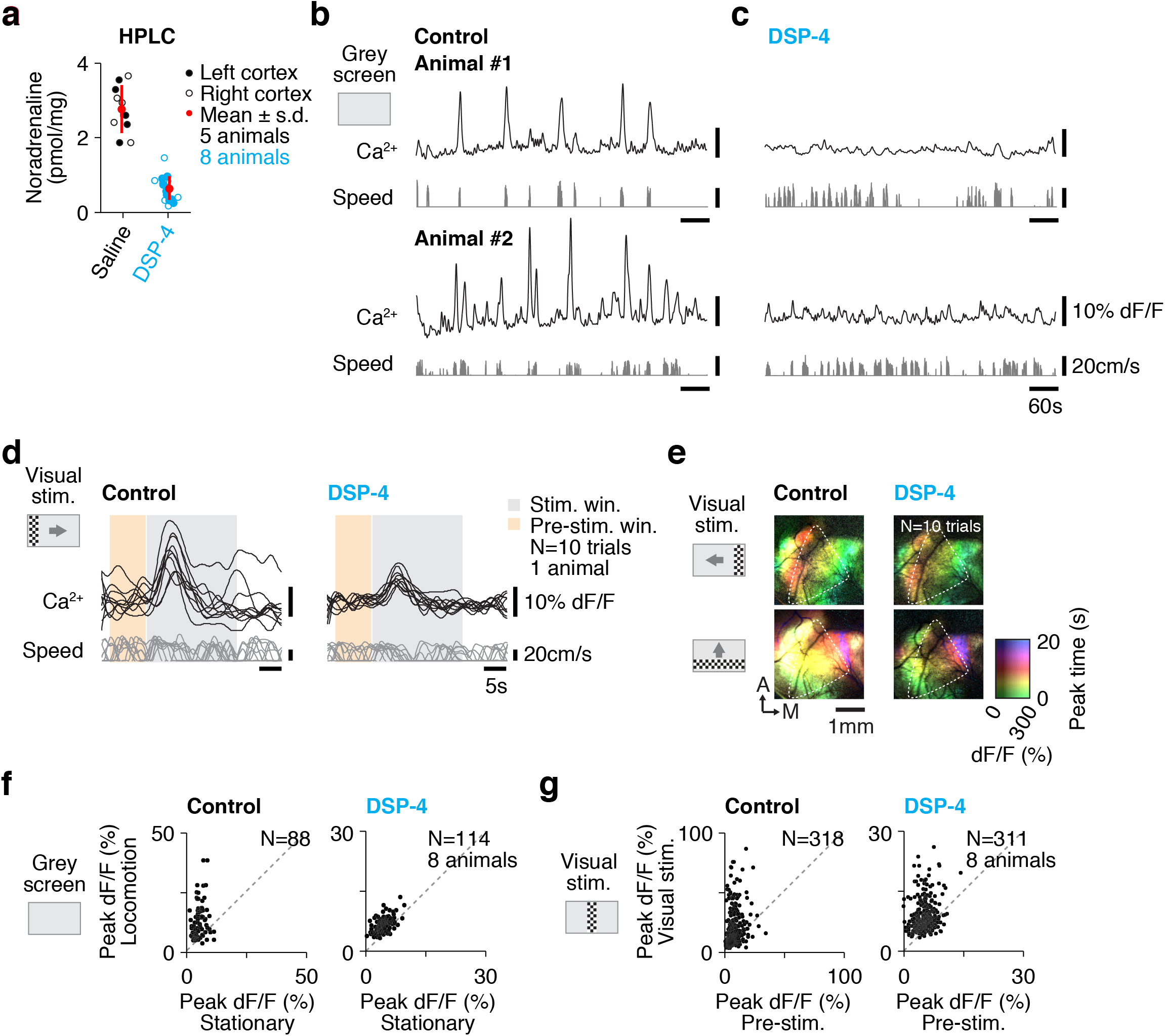
Visually-induced and behavioral state-induced signals operate through distinct mechanisms. (a) Scatter plot of noradrenaline concentration in bulk extracts from left (imaged) cortex and right cortex. Noradrenaline was measured by HPLC seventy-two hours after intraperitoneal injection of DSP-4. Control mice were injected with saline. Red bars indicate mean ± s.d. (b, c) Noradrenaline depletion abolishes the Ca^2+^ responses associated with locomotion onset. Ca^2+^ transients (top traces) recorded from two exemplar mice presented with a grey screen during spontaneous locomotion (bottom traces) before (b) and after (c) systemic DSP-4 administration. (d, e) Visually-induced responses are partially preserved after depletion of noradrenaline. Example Ca^2+^ transients (top traces) from an animal before (left) and after (right) DSP-4 administration. Visual stimuli (grey box) were presented moving in the indicated directions during spontaneous locomotion (bottom traces) (d). Pseudo-color, pixel-wise maps of trial averages for vision-induced increases in GCaMP6 fluorescence. Maps are the average of 10 trials per stimulus direction in 1 animal before and after DSP-4 administration (e). (f, g) DSP-4 differentially affects the responses induced by visual stimulation or locomotor activity. Scatter plots showing the peak Ca^2+^ activity for locomotion onset-induced (f) and visually-induced transients (g) before (left) and after (right) DSP-4 administration. Dashed lines indicate unity.

Together, these data suggest that co-incident detection of sensory inputs and behavioral state occurs at the level of individual astrocytes, and is not due to the combination of locomotion and visual stimuli leading to more firing of LC neurons, that release noradrenaline in the visual cortex^17^. In our model, noradrenaline acts to directly gate the visual response in astrocytes, which is entirely consistent with reports of enforced locomotion enhancing astrocyte response to a simple visual stimulus (uniform light flash)^7^. It also reconciles earlier reports where responses to visual stimuli were not detected or weak, but which did not take into account the influence of behavior^10^.

Our results stand in contrast to those observed in the ferret^6^, where synaptic activity alone was sufficient to elicit retinotopic astrocyte responses. This may reflect differences between astrocyte function in lightly anesthetized and awake behaving animals. Alternatively, it may reflect the differing organization of the visual cortex between the two species: due to the columnar organization of the cortex in ferret, the majority of neurons within an astrocyte’s territory will fire synchronously when presented with a grating in a particular orientation. In contrast, orientation preference is arranged in a salt-and-pepper fashion in mouse cortex, meaning effective responses will only occur in a fraction of the neighboring neurons. Furthermore, firing rates evoked by a grating stimulus are nearly three times higher in ferret than in mouse visual cortex^11^. Hence, astrocytes in mouse cortex receive comparatively sparse input, which might require cells to be ‘primed’ prior to synaptic activity.

In summary, we show that cortical astrocytes integrate signals across input modalities, with behavioral state directly influencing sensory responses. Such a system is likely to underlie many aspects of astrocyte function regulated by fluctuations in intracellular Ca^2+^, for example visual synaptic plasticity^18^. Our data support a role for astrocytes in translating the brain-wide action of neuromodulators to produce specific functional effects on local neuronal circuits^17, 19^. Greater understanding of how such systems operate will undoubtedly further our understanding of co-ordinated CNS function^20^.

## Material and Methods

Detailed methods are available as Supplementary Information.

## Acknowledgements

MGH gratefully acknowledges support from the European Research Council (Starting Grant 281961), Fonds voor Wetenschappelijk Onderzoek Vlaanderen (FWO) (Grants 1523014N and 1527315N) and VIB Institutional Funding. MS was supported by a Marie Curie Intra-European Fellowship (331018). VB acknowledges support from FWO (Grant G0D0516N), KU Leuven Research Council (Grant C14/16/048) and NERF Institutional Funding. NERF is funded by Imec, VIB and KU Leuven. We thank João Couto for providing code for pupil detection, Filip de Vin for technical assistance during experiments, and Benjamin Puccio, Molly Kirk, Jessica Mitchell, Daniel Tovbin, Alicja Ronisz and Frederique Ooms for help with animal training. We thank Dr. Jérôme Wahis for critical comments on the manuscript.

## Author Contributions

MS, SK, VB and MGH devised the research; MS and SK designed the experiments; MS performed the experiments; SK and MS analyzed the data; PPVV performed HPLC-based neurotransmitter analyses; MS, SK, VB and MGH wrote the manuscript.

